# Accessible Analysis of Longitudinal Data with Linear Mixed Effects Models

**DOI:** 10.1101/2020.12.03.411058

**Authors:** Jessica I. Murphy, Nicholas E. Weaver, Audrey E. Hendricks

**Affiliations:** Mathematical and Statistical Sciences, University of Colorado Denver; Biostatistics and Informatics, Colorado School of Public Health

**Keywords:** ANOVA, linear mixed effects, longitudinal, microbiome, mouse, Shiny app

## Abstract

Longitudinal mouse models are commonly used to study possible causal factors associated with human health and disease. However, the statistical models, such as two-way ANOVA, often applied in these studies do not appropriately model the experimental design, resulting in biased and imprecise results. Here, we describe the linear mixed effects (LME) model and how to use it for longitudinal mice studies. We re-analyze a dataset published by Blanton et al (Science 2016) that modeled mice growth trajectories after microbiome implantation from nourished or malnourished children. We compare the fit and stability of different parameterizations of ANOVA and LME models; most models found the nourished vs. malnourished growth trajectories differed significantly. We show through simulation that the results from the two-way ANOVA and LME models are not always consistent. Incorrectly modeling correlated data can result in increased rates of false positives or false negatives, supporting the need to model correlated data correctly. We provide an interactive Shiny App to enable accessible and appropriate analysis of longitudinal data using LME models.

## INTRODUCTION

Longitudinal mouse models are often used in biomedical research to better understand human conditions. Common applications include studying the effects of the gut microbiome on human health and disease (Blanton et al., 2016; Britton et al., 2019; Feehley et al., 2019; Ridaura et al., 2013; Tanoue et al., 2019) as well as studying working and long-term memory in Alzheimer’s disease (Alamed et al., 2006; Cracchiolo et al., 2007; Gajbhiye et al., 2017; Rosenzweig et al., 2019). In these studies, mice are randomly assigned to a treatment group and a continuous outcome, such as growth, is tracked across time. The goal of longitudinal mouse models is usually to determine whether the trajectories of mice over time vary by treatment group.

Correlation between observational units (i.e. units by which the data is gathered) is inherent within longitudinal mouse models due to the longitudinal and sometimes nested study design. For instance, longitudinal measurements taken from the same mouse are more likely to be similar than measurements taken from different mice. Experiments sometimes also have a nested design where the fecal sample from a single donor is transplanted into multiple mice (Blanton et al., 2016; Britton et al., 2019; Feehley et al., 2019). Mice with transplanted microbiota from the same donor are likely to have more similar (i.e. correlated) microbiota profiles. Incorrectly modeled experimental structure can result in biased and imprecise estimates, which can lead to inaccurate conclusions (Cheng et al., 2010; Verbeke and Molenberghs, 2000).

Correctly modeling correlated data requires careful consideration. In many longitudinal mouse studies, a two-way ANOVA model is used (Blanton et al., 2016; Ridaura et al., 2013; Tanoue et al., 2019), with factors (i.e. predictor variables) for treatment group, time, and the interaction between treatment group and time. A two-way ANOVA assumes independent observations and thus does not account for correlation from longitudinal or nested measurements. Repeated measures ANOVA is also commonly used (Cracchiolo et al., 2007; Rosenzweig et al., 2019), which does control for the correlation of measurements due to longitudinal structure but not nested designs. The linear mixed effects (LME) model is a flexible method enabling correct modeling of both longitudinal and nested correlation (Feehley et al., 2019).

Here, we describe the LME model and how to use it for longitudinal mice studies. We also re-analyze a dataset from Blanton et al. published in Science in 2016 that investigated the growth trajectories of mice from nourished and malnourished children (Blanton et al., 2016). Finally, we provide an interactive Shiny App so others can easily implement an appropriate statistical analysis for longitudinal mice studies.

## RESULTS

### Linear Mixed Effects Models

A traditional linear model is defined by the following formula,

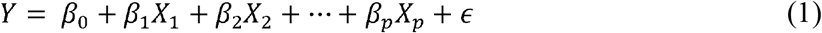

*where Y is the outcome variable, X*_l_,…, *X*_*p*_ *are the predictors or factors, β*_0_, … , *β*_*p*_ *are the regression parameters reflecting the relationship between each X*_*j*_ *and Y while controlling for the other predictors, and ϵ is the random error.*

When used for longitudinal models, trajectories by group are often modelled using formula 2 below. This is identical to a two-factor ANOVA with interaction. The group by time interaction is usually the effect of interest in longitudinal studies and identifies whether trajectories differ by group.

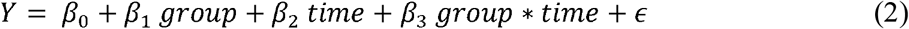

*where group refers to the treatment group, time refers to the time measurement (continuous), and group*\**time refers to the interaction between treatment group and time. The model parameters are defined in **Table 1**.*

**Table 1:** Parameter Definitions

| Parameter | Definition |
| --- | --- |
| $\beta_0$ | Intercept |
| $\beta_1$ | Effect of treatment group (e.g. undernourished vs healthy)* |
| $\beta_2$ | Effect of time* |
| $\beta_3$ | Effect of the treatment group by time interaction |
| $\gamma_{D0}$ | Random intercept for donor |
| $\gamma_{D1}$ | Random slope for donor |
| $\gamma_{M0}$ | Random intercept for mouse nested within donor |
| $\gamma_{M1}$ | Random slope for mouse nested within donor |
| $\epsilon$ | Random error |

\*The effect of time and the effect of group should not be analyzed or interpreted individually given the interaction term.

The random error term, *ϵ*, captures variation in the outcome, *Y*, not explained by the model. In a linear model, the random error is assumed to be independent and identically distributed with a constant variance. The independence assumption is violated if unaccounted correlation is present, which occurs in longitudinal data and nested designs.

To account for correlation between observations, LMEs use random effects to explicitly model the correlation structure, thus removing correlation from the error term. Random effects can take many functional forms (e.g. linear, quadratic) depending on the structure of the data. Here we focus on two commonly used random effects: a random intercept and a random slope. For example, a random intercept for mouse allows the mean value (i.e. intercept) of *Y* to differ between mice while a random slope term allows the rate of change (i.e. slope) between a predictor such as time and the outcome to differ between mice. A random slope in addition to a random intercept allows both the rate of change and the mean value to vary by mouse. Random slopes are usually only used with random intercepts.

A random slope, *γ*_*i*1_, and intercept, *γ*_*i*0_, model for mouse *i* is shown in equation 3 below.

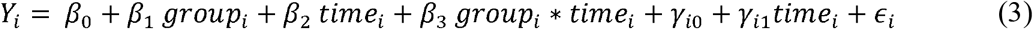

In this model, *β*_0_, *β*_1_, *β*_2_, and *β*_3_ are often referred to as fixed effects because they are the same for all subjects. Random effects are usually specified for the repeated unit. Here, the repeated units are mouse (multiple time points per mouse) and donor (multiple mice nested within a single donor). The mouse would be considered the nested random effect and the donor would be considered the clustered random effect.

When in doubt or if the structure of the data is unknown, the full model, which includes a slope and intercept for each random effect, is commonly used (Cheng et al., 2010; Verbeke and Molenberghs, 2000). Then, the need for each slope and/or intercept can be tested for inclusion in the model as further described.

### Reanalysis of Blanton et. al Dataset

Five models were evaluated in the reanalysis of the Blanton et. al dataset. The models are described in the following section, and the model parameters are defined in **Table 1**. The ***Full Model*** (Model I) is specified below, where *Y* refers to the outcome variable (growth), *group* refers to the treatment group (undernourished vs healthy), *time* refers to the number of days, and *group*\**time* refers to the interaction between treatment group and time. The ***Nested Models*** (Models II – V) include a subset of terms from the ***Full Model***.

### Full Model

**Donor Intercept & Slope + Mouse Intercept & Slope Random Effects:**

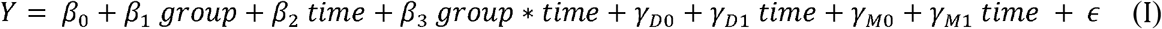

### Nested Models

**Donor Intercept + Mouse Intercept & Slope Random Effects:**

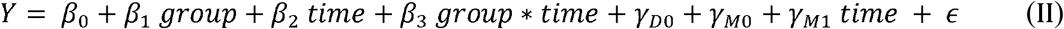

**Mouse Intercept & Slope Random Effects:**

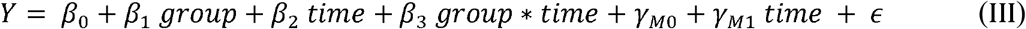

**Mouse Intercept Random Effect:**

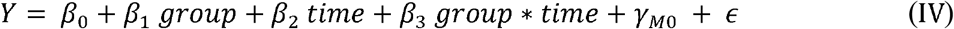

**No Random Effects:**

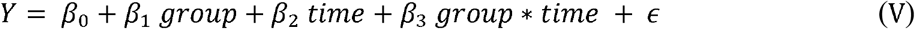

Hypothesis tests of the random effect terms (i.e. tests of nested models) were performed with a likelihood ratio test (LRT) as well as a parametric bootstrap test. For both, each additional random effect term significantly improved model fit indicating that the full model (Model I) is preferred (**Table 2**). However, Model I produced convergence warnings for three out of seven optimizers checked (bobyqa, Nelder_Mead, and optimx.L-BFGS-B), suggesting Model I is less stable. Model II was significantly better than Model III and did not produce any convergence warnings.

**Table 2:** Assessment of Nested Models

| Models | Likelihood Ratio Test |  | Parametric Bootstrap Test |  |
| --- | --- | --- | --- | --- |
|  | Test Statistic | P-value | Test Statistic | P-value |
| I vs II | 9.85 | 0.00727 | 9.62 | 0.00102 |
| II vs III | 7.99 | 0.00471 | 7.72 | 0.00193 |
| III vs IV | 419 | < 2.2e-16 | 418 | 0.00100 |
| IV vs V | 243 | < 2.2e-16 | ---* | --- |

\*The parametric bootstrap test was unable to be used to compare models IV and V because the R function *PBmodcomp* only compares models of the same type. Model IV is a mixed effects model whereas model V is a linear model.

The interaction effect estimate is smaller for Model V compared to Models I through IV for which the estimates are fairly consistent. The standard error estimates for the interaction effect vary more between models. The standard error estimate of the interaction term is largest for Model I, smaller for Models II and III, and even smaller for Models IV and V (**Table 3**).

**Table 3:** Model Comparison

| Model | Interaction Effect |  |  | Log Likelihood |
| --- | --- | --- | --- | --- |
| | $\hat{\beta}$ | SE | p-value | |
| I | -0.42 | 0.26 | 0.15 | -1,178.74 |
| II | **-0.42** | 0.14 | 0.0058 | -1,184.60 |
| III | **-0.43** | 0.14 | 0.0051 | -1,189.72 |
| IV | **-0.40** | 0.05 | 1.88e-14 | -1,399.33 |
| V | **-0.32** | 0.07 | 1.48e-5 | -1,518.64 |

\*Significant interaction terms ( $p < 0.05$ ) are bolded.

**Table 4:** Type I Error and Power

|  | Mouse Intercept & Slope<br>Simulation Scenario |  |  | Mouse Intercept<br>Simulation Scenario |  |  | No Random Effects<br>Simulation Scenario |  |  |
| --- | --- | --- | --- | --- | --- | --- | --- | --- | --- |
| Effect<br>Size | Mouse<br>Intercept<br>& Slope<br>(Model III) | Mouse<br>Intercept<br>(Model IV) | No Random<br>Effects<br>(Model V) | Mouse<br>Intercept<br>& Slope<br>(Model III) | Mouse<br>Intercept<br>(Model IV) | No Random<br>Effects<br>(Model V) | Mouse<br>Intercept<br>& Slope<br>(Model III) | Mouse<br>Intercept<br>(Model IV) | No Random<br>Effects<br>(Model V) |
| 0%* | 0.040<br>[0.036, 0.044] | 0.334<br>[0.325, 0.343] | 0.314<br>[0.305, 0.323] | 0.011<br>[0.009, 0.013] | 0.019<br>[0.016, 0.022] | 0.017<br>[0.014, 0.019] | 0.028<br>[0.025, 0.031] | 0.050<br>[0.045, 0.054] | 0.049<br>[0.045, 0.054] |
| 10% | 0.05<br>[0.03, 0.06] | -.** | - | 0.19<br>[0.16, 0.21] | 0.26<br>[0.24, 0.29] | 0.24<br>[0.22, 0.27] | 0.26<br>[0.23, 0.29] | 0.40<br>[0.37, 0.43] | 0.40<br>[0.37, 0.43] |
| 25% | 0.08<br>[0.06, 0.1] | - | - | 0.92<br>[0.90, 0.93] | 0.98<br>[1.00, 1.00] | 0.97<br>[0.96, 0.98] | 0.92<br>[0.90, 0.94] | 0.99<br>[0.98, 0.99] | 0.99<br>[0.98, 0.99] |
| 50% | 0.23<br>[0.21, 0.26] | - | - | 1.00<br>[0.99, 1.00] | 1.00<br>[1.00, 1.00] | 1.00<br>[1.00, 1.00] | 1.00<br>[1.00, 1.00] | 1.00<br>[1.00, 1.00] | 1.00<br>[1.00, 1.00] |
| 100% | 0.72<br>[0.69, 0.74] | - | - | 1.00<br>[1.00, 1.00] | 1.00<br>[1.00, 1.00] | 1.00<br>[1.00, 1.00] | 1.00<br>[1.00, 1.00] | 1.00<br>[1.00, 1.00] | 1.00<br>[1.00, 1.00] |

\*Type I Error
\*\*Power is only compared when Type I Error was close to or below the expected level of 0.05.

### Simulations

To compare the type I error and power of the models, data was simulated for three different covariance scenarios: a random intercept and slope for mouse (Model III), a random intercept for mouse (Model IV), and no random effects (Model V).

Across all simulation scenarios, the model that matches the simulation scenario performs best by maintaining the type I error and having higher power. For instance, for the mouse intercept and slope simulation scenario, the model that matches the simulation scenario (Model III) is the only model that maintains the appropriate type I error. The type I error is severely inflated for Models IV and V, which do not include the slope random effect term. For the mouse intercept simulation scenario, Model IV, which matches the simulation scenario, has slightly more power than Models III and V. But, the type I error is very conservative for all three models, even the model that matches the simulation scenario (Model IV). In supplemental **Table S1** with N = 40 independent mice, the increased power for Model IV is better highlighted as the type I error is no longer conservative for this larger sample size. For the simulation scenario with no random effects, Model V, which matches the simulation scenario, and Model IV are both close to the expected value of 0.05, while Model III has a slightly conservative type I error estimate. Consequently, Model III also has slightly less power than Models IV and V.

To determine if the small simulated sample size of 8 donors with 5 mice nested within each donor was affecting the type I error estimates, a simulation analysis was performed for larger sample sizes (**Table 5**). While the nested model of five mice per eight donors results in 40 mice, mice within the same donor are correlated (i.e. contain dependent information), resulting in an effective independent sample size less than 40. Conversely, the scenario with 40 donors, one for each mouse, provides independent information for each mouse. Indeed, for the mouse intercept simulation scenario, the type I error for Model IV is deflated for the five mice per eight donors sample size, but is close to the expected value of 0.05 when there are 40 independent mice. For the mouse intercept simulation scenario, Models III and V, which do not match the experimental design of the simulation scenario, have slightly and considerably deflated type I error, respectively. Model III, which contains an unnecessary random slope term, approaches the expected type I error as the sample size increases (**Table 5**). Model V, which does not include the necessary random intercept term, has severely deflated type I error across all sample sizes. For the mouse intercept and slope simulation scenario, the type I errors for Models IV and V are severely inflated across all sample sizes.

**Table 5:**
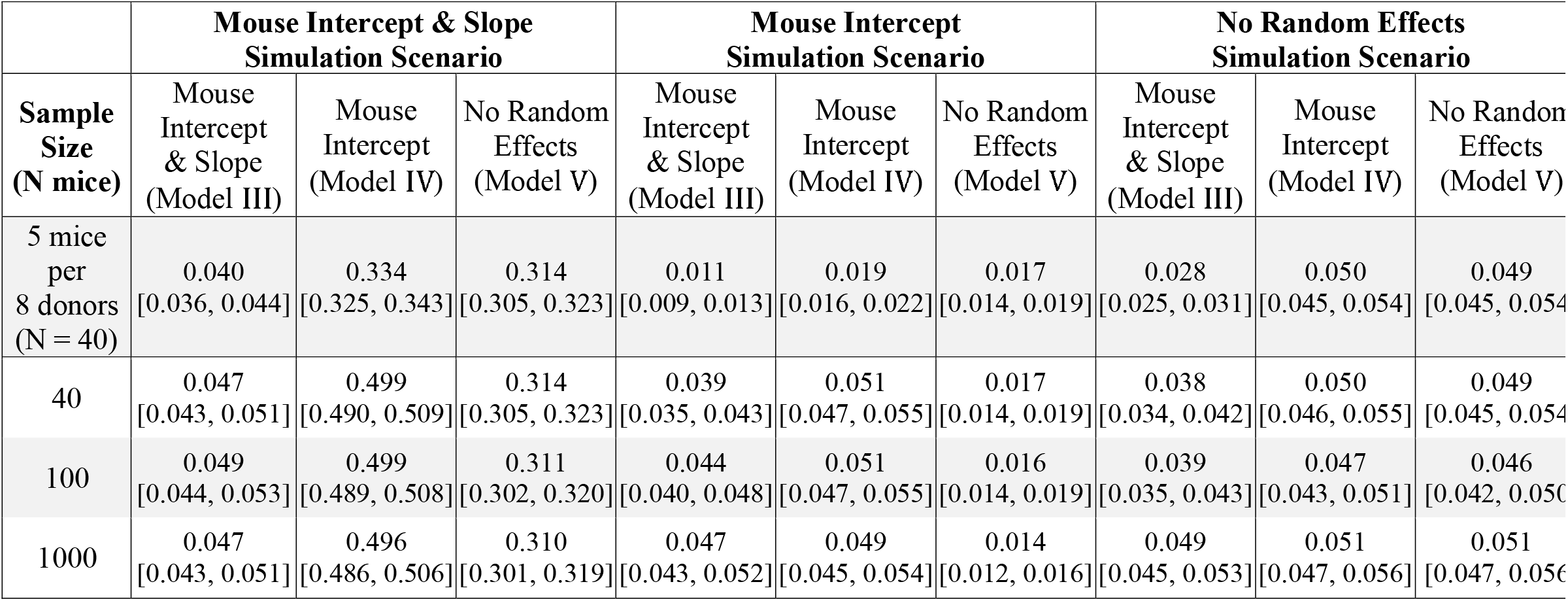
Type I Error by Sample Size

| **Sample Size**<br>**(N mice)** | **Mouse Intercept & Slope**<br>**Simulation Scenario** |  |  | **Mouse Intercept**<br>**Simulation Scenario** |  |  | **No Random Effects**<br>**Simulation Scenario** |  |  |
| --- | --- | --- | --- | --- | --- | --- | --- | --- | --- |
|  | Mouse Intercept & Slope (Model III) | Mouse Intercept (Model IV) | No Random Effects (Model V) | Mouse Intercept & Slope (Model III) | Mouse Intercept (Model IV) | No Random Effects (Model V) | Mouse Intercept & Slope (Model III) | Mouse Intercept (Model IV) | No Random Effects (Model V) |
| 5 mice per 8 donors (N = 40) | 0.040<br>[0.036, 0.044] | 0.334<br>[0.325, 0.343] | 0.314<br>[0.305, 0.323] | 0.011<br>[0.009, 0.013] | 0.019<br>[0.016, 0.022] | 0.017<br>[0.014, 0.019] | 0.028<br>[0.025, 0.031] | 0.050<br>[0.045, 0.054] | 0.049<br>[0.045, 0.054] |
| 40 | 0.047<br>[0.043, 0.051] | 0.499<br>[0.490, 0.509] | 0.314<br>[0.305, 0.323] | 0.039<br>[0.035, 0.043] | 0.051<br>[0.047, 0.055] | 0.017<br>[0.014, 0.019] | 0.038<br>[0.034, 0.042] | 0.050<br>[0.046, 0.055] | 0.049<br>[0.045, 0.054] |
| 100 | 0.049<br>[0.044, 0.053] | 0.499<br>[0.489, 0.508] | 0.311<br>[0.302, 0.320] | 0.044<br>[0.040, 0.048] | 0.051<br>[0.047, 0.055] | 0.016<br>[0.014, 0.019] | 0.039<br>[0.035, 0.043] | 0.047<br>[0.043, 0.051] | 0.046<br>[0.042, 0.050] |
| 1000 | 0.047<br>[0.043, 0.051] | 0.496<br>[0.486, 0.506] | 0.310<br>[0.301, 0.319] | 0.047<br>[0.043, 0.052] | 0.049<br>[0.045, 0.054] | 0.014<br>[0.012, 0.016] | 0.049<br>[0.045, 0.053] | 0.051<br>[0.047, 0.056] | 0.051<br>[0.047, 0.056] |

These results suggest that leaving out a necessary random effect term will result in inaccurate type I error even at large sample sizes. This is seen with Models IV and V for the mouse intercept and slope simulation scenario and with Model V for the mouse intercept simulation scenario. Importantly, the direction of the bias in the type I error can result in too conservative (i.e. too small type I error) or anti-conservative (i.e. too large type I error) results. Models that include an unnecessary random effect term, e.g. Model III in the intercept only simulation scenario or Models III or IV in the no random effects simulation scenario, can result in deflated type I error at smaller sample sizes. However, this bias decreases as the sample size increases ultimately resulting in the appropriate type I error. These results suggest that the type I error is better controlled when random effect terms are included but may result in lower power as shown in **Table 2**. Regardless, as we discuss above and show in **Table 2**, whether including random effect terms results in better model fit can be tested.

### EasyLME: A Shiny App

Since analyzing longitudinal mice data requires careful consideration and the use of LME models is not always easily available, we provide an interactive and user-friendly Shiny App that allows others to implement and choose an appropriate LME model more easily (Chang et al., 2020). The app, called EasyLME, can be accessed through any web browser at https://jessmurphy.shinyapps.io/EasyLME/. To explore the app’s features, users can upload their own data or use the demo data from Blanton et al. The app assumes the response and time variables are continuous and the grouping variable as well as the random effects are factors. Users can then select variables representing the structure of their data including an option to specify if the model contains nested random effects. Users can also select additional covariates, if desired. For the demo data, the random effect variables are “Mouse” nested in “Donor” as shown on the left-hand side of **Figure 1**, since samples from each donor were transplanted into multiple mice. Once the variables are chosen, the user can navigate through the output tabs to see the various aspects of the analysis. This basic design of the app was inspired by Medplot, a Shiny app for longitudinal medical data (Ahlin et al., 2015).

**Figure 1:**
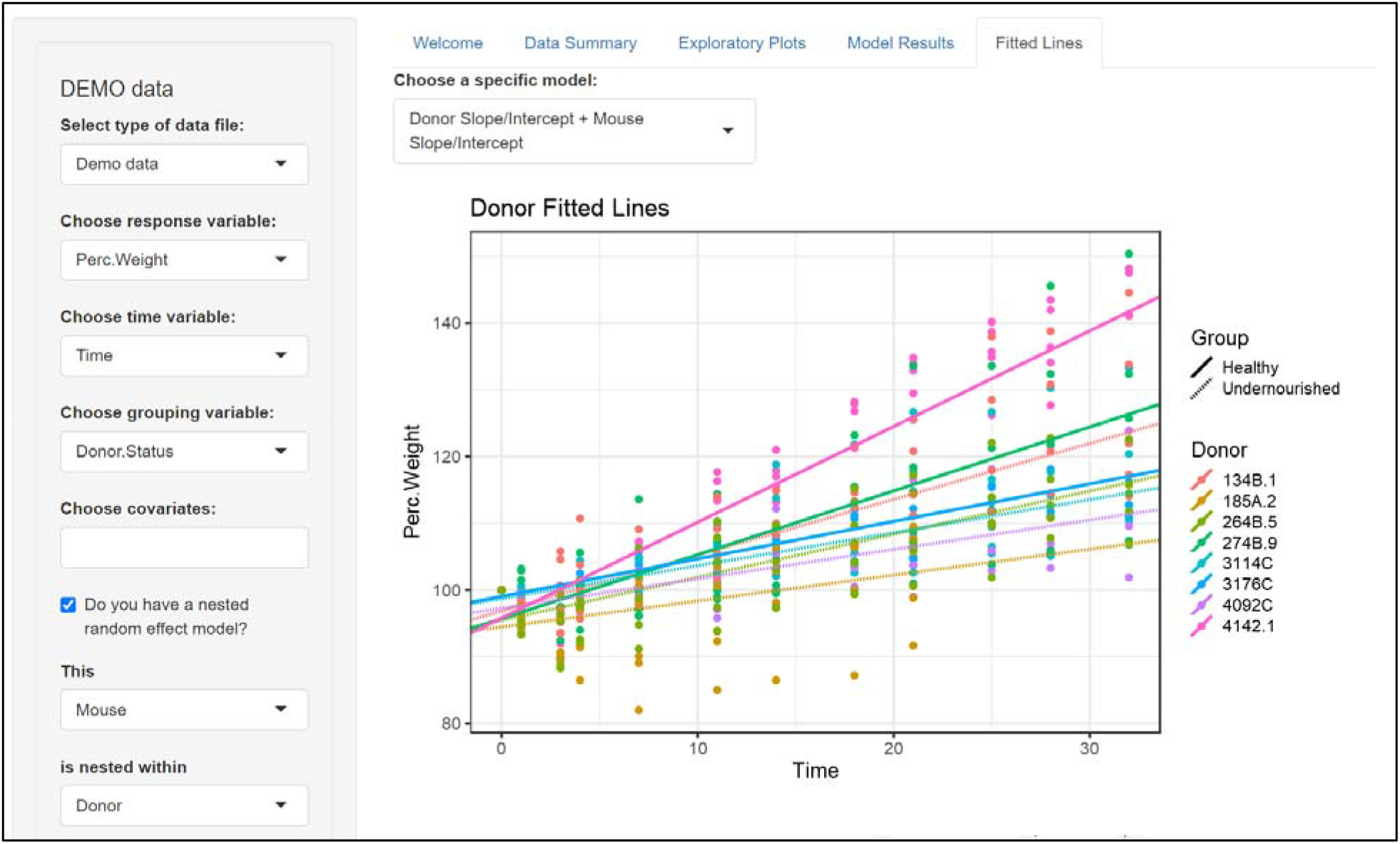
Shiny App User Interface

The app contains five tabs as shown in **Figure 1**: *Welcome*, *Data Summary*, *Exploratory Plots*, *Model Results*, and *Fitted Lines*. The main page of the app is the ***Welcome*** tab with an *About* section explaining the app functions followed by a *Getting Started* section. The ***Data Summary*** tab provides univariate summaries of each variable selected by the user. The data summaries help ensure that the appropriate variables are selected and can highlight any extreme or missing values.

The ***Exploratory Plots*** tab provides two visualizations helpful to understand structure in the data. The *Overall Trends* scatterplot (**Figure S1**) visualizes the relationship between the response variable and time. This plot is useful for checking the assumption of linearity between the response and time. The *Donor Trendlines* plot (**Figure S2**) displays the average response over time for the clustered random effect (e.g. donor). This plot helps to visualize if a random intercept and/or random slope would be appropriate for the clustered random effect. It also shows the presence of missing data if there are gaps in the lines or the lines stop short. If the data are not nested, this graph will just show the trendlines for the random effect without the need for averaging.

The ***Model Results*** tab contains a comparison table for testing nested models. The table contains coefficient estimates with standard errors in parentheses, log likelihoods, and p-values from likelihood ratio tests of the nested models. These results are similar to those of **Table 2** and **Table 3**. If the data are nested (e.g. multiple mice per donor), the table compares the five models listed above in the ***Reanalysis of Blanton et. al Dataset*** section. If the data are not nested (e.g. one mouse per donor), the table only compares Models III - V (mouse intercept and slope, mouse intercept, and no random effects). Below the table, the user can select a specific LME model from the drop-down menu to see more detailed information, such as the random effect estimates.

Lastly given a selected model, the ***Fitted Lines*** tab visualizes the fitted line for each unit (e.g. donor or mouse) as seen in **Figure 1.** If the data are nested, the tab will contain two plots, one for each level (e.g. mouse and donor). These plots are useful for visualizing how the inclusion of a random slope and intercept versus just a random intercept affects the model fit. The inclusion of a random intercept versus no random effects can also be compared.

## DISCUSSION

Longitudinal data, and especially nested longitudinal data, are often modelled incorrectly. ANOVA, although commonly used, does not accurately account for correlation from repeated measurements or nested structure. Here, we compare the LME model with the ANOVA in a gut microbiome case study involving longitudinal mouse models. In our reanalysis of Blanton et al., we find evidence that LME models based on the experimental design better fit the data. Importantly, we show that results from the LME model and ANOVA will not always produce consistent results in terms of effect estimate, standard error, and subsequently statistical significance. Through simulations, we show that the type I error is not well controlled when using an ANOVA model for longitudinal data and can result in false positives or false negatives. We also show that LMEs, with the inclusion of necessary random effect terms to appropriately model the experimental structure, produce unbiased type I error at sufficiently large sample sizes.

To allow easier implementation of LME models, we designed EasyLME, a Shiny app to enable appropriate analysis of longitudinal mice studies. The app’s easy-to-use features, such as the exploratory plots, model tests/comparisons, and fitted lines, allow researchers to choose the most suitable model for their data. Although the app was designed for mouse studies, it is applicable to longitudinal studies to identify group differences in trajectory for a continuous outcome.

While the goal of the app is to make analyzing longitudinal data with LMEs more accessible, consultation with a statistician is still recommended in study design, analysis, and interpretation of results. We also recommend that, prior to analysis, users create a data analysis plan that describes quality control, significance thresholds, and how the final model will be chosen (Cheng et al., 2010; Michener, 2015; Simpson, 2015). Additionally, we advise the user to consider both model fit and stability when selecting a model (e.g. visualizations, nested model comparisons, as well as convergence warnings). If the results and conclusions vary greatly between the models, such as large changes in the effect size or significance of the interaction term, we recommend reporting the results from all models considered. Inconsistent results between the models would suggest there is likely additional uncertainty in the outcome not explained by the current model structure, which would require further investigation.

Although LME models can be performed in other programs (e.g. STATA, SAS, SPSS), R was used for this analysis since it is a widely known, free, and open source platform. As such, we hope our app will increase accessibility for applied researchers working with longitudinal data.

## MATERIALS and METHODS

### Blanton et al. Dataset

The dataset used in this paper was obtained from a 2016 publication titled “Gut bacteria that rescue growth impairments transmitted by immature microbiota from undernourished children” (Blanton et al., 2016). In the experiment of interest, fecal samples from eight healthy and eleven undernourished children aged 6 or 18 months were obtained. Each fecal sample was orally transferred to five germ-free mice. The percent weight change of each mouse was recorded at 12 time points: 0, 1, 3, 4, 7, 11, 14, 18, 21, 25, 28, and 32 days. In the original study, the statistical analysis was restricted to eight donor samples (three healthy, five underweight) that produced >50% transplantation efficiency resulting in a total of 480 observations across all time points, mice, and donors. Forty-three observations were missing resulting in 437 observations. The study design for this dataset is shown in **Figure 2**.

**Figure 2:**
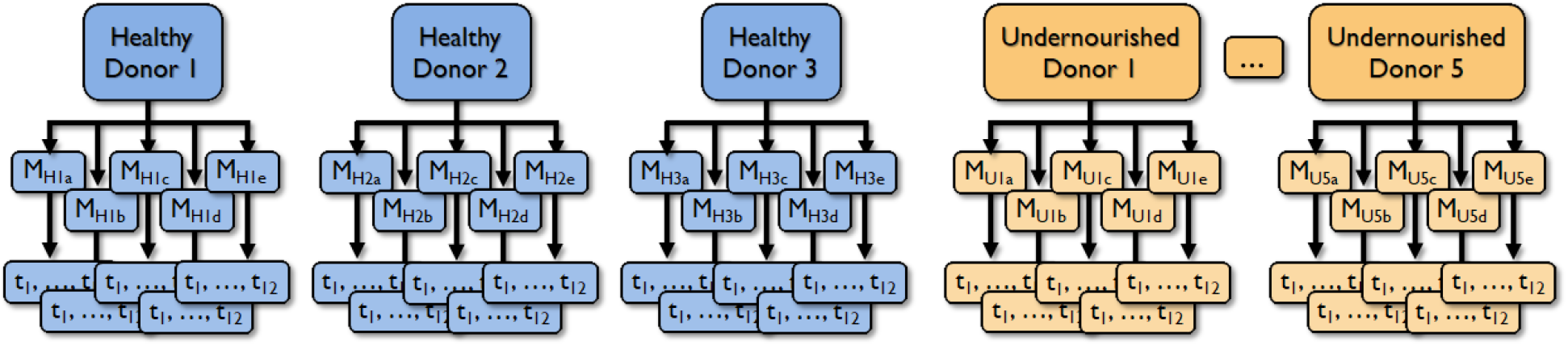
Study Design where M refers to an individual mouse nested within a donor, the numbered subscripts H and U refer to healthy and undernourished donors, respectively, and t refers to an individual time point nested within a mouse.

### Reanalysis of Blanton et. al Dataset

Analysis of the Blanton et. al data with LME models was performed using the *lme4* v1.1-23 and *lmerTEST* v3.1-2 packages in R v4.0.3 (Bates et al., 2015; Kuznetsova et al., 2017; R Core Team, 2020). The coefficients were estimated using restricted maximum likelihood (REML) (Verbeke and Molenberghs, 2000) and the random effects terms were allowed to be correlated. The fit of each model was measured using log likelihoods, with a larger log likelihood indicating a better fit. The stability of the model estimates for different optimizers was checked using the *allFit* function from the *lme4* v1.1-23 R package (Bates et al., 2015). The estimates were considered stable if they did not change substantially across the optimizers and if the selected optimizer did not produce any convergence warnings.

The LRT is an appropriate method for hypothesis testing of fixed effects terms. However, the LRT can produce inaccurate p-values for random effects since random effect estimates are tested against zero, which is on the boundary of the parameter space (variance must be greater than or equal to zero) (Verbeke and Molenberghs, 2000). As such, parametric bootstrap methods (*PBmodcomp* function from the *pbkrtest* package) in addition to LRTs were used to perform hypothesis tests of random effect terms (Halekoh and Højsgaard, 2014). The *PBmodcomp* function can only be used to compare models of the same type and thus could not be used to test a LME model (Model IV) vs a linear model (Model V). When completing hypothesis tests of nested models, each model was refit using maximum likelihood (ML) estimation. For all statistical tests, a significance threshold of p < 0.05 was used.

### Simulations

Data was simulated from a multivariate normal distribution (*mvrnorm* function from the *MASS* package) for three different covariance scenarios: a random slope and intercept for mouse (Model III), a random intercept for mouse (Model IV), and no random effects (Model V) (Venables and Ripley, 2002). The simulation sample size consisted of four healthy donors and four undernourished donors, with five mice per donor. The same twelve time points used in Blanton et al. were used for simulations: 0, 1, 3, 4, 7, 11, 14, 18, 21, 25, 28, and 32 days. The simulation parameters for the fixed effect estimates for treatment group, time, and time by treatment interaction from Model III were used for all simulation scenarios because this model produced the largest interaction effect. The interaction effect between treatment group and time was varied to be 0, 10, 25, 50, and 100% of the observed effect from Model III for the Blanton et al. data, while keeping the simulation parameters for the group and time effect estimates the same. The mice were assumed to be independent and the same variance matrix was assumed for all mice within a simulation scenario.

LME models III, IV, and V were fit to the simulated data using the default optimizer within *lme4*. Type I error and power were calculated using 10,000 and 1,000 simulation replicates, respectively. Type I error was assessed for an interaction effect of 0. Since the asymptotic properties of LME models may not be reached for small sample sizes, type I error was also evaluated at larger sample sizes (N = 40, 100, 1000), with one mouse per donor and an equal number of donors per treatment group. All hypothesis tests were assessed at α = 0.05.

The original dataset and the R code for these analyses can be found at https://github.com/JessMurphy/Longitudinal.Mouse.

## ACKNOWLEDGEMENTS

We appreciate Pitshou Nzazi Duki for helping design a preliminary version of the Shiny app and Drs. Minghua Tang and Genevieve Roberts for beta testing the app.

## DATA AVAILABILITY

The original dataset and the R code for the analyses can be found at https://github.com/JessMurphy/Longitudinal.Mouse while the Shiny app R code can be found at https://github.com/JessMurphy/EasyLME. The Shiny app can be accessed through any web browser at https://jessmurphy.shinyapps.io/EasyLME/.

## REFERENCES

Ahlin, Č., Stupica, D., Strle, F. and Lusa, L. (2015). medplot: a web application for dynamic summary and analysis of longitudinal medical data based on R. PLoS One 10, e0121760.

Alamed, J., Wilcock, D. M., Diamond, D. M., Gordon, M. N. and Morgan, D. (2006). Two-day radial-arm water maze learning and memory task; robust resolution of amyloid-related memory deficits in transgenic mice. Nature protocols 1, 1671.

Bates, D., Mächler, M., Bolker, B. and Walker, S. (2015). Fitting linear mixed-effects models using lme4. Journal of Statistical Software 67, 1–48.

Blanton, L. V., Charbonneau, M. R., Salih, T., Barratt, M. J., Venkatesh, S., Ilkaveya, O., Subramanian, S., Manary, M. J., Trehan, I., Jorgensen, J. M. et al. (2016). Gut bacteria that prevent growth impairments transmitted by microbiota from malnourished children. Science 351.

Britton, G. J., Contijoch, E. J., Mogno, I., Vennaro, O. H., Llewellyn, S. R., Ng, R., Li, Z., Mortha, A., Merad, M. and Das, A. (2019). Microbiotas from humans with inflammatory bowel disease alter the balance of gut Th17 and RORγt+ regulatory T cells and exacerbate colitis in mice. Immunity 50, 212–224. e4.

Chang, W., Cheng, J., Allaire, J., Xie, Y. and McPherson, J. (2020). shiny: Web Application Framework for R.

Cheng, J., Edwards, L. J., Maldonado◻Molina, M. M., Komro, K. A. and Muller, K. E. (2010). Real longitudinal data analysis for real people: building a good enough mixed model. Statistics in medicine 29, 504–520.

Cracchiolo, J. R., Mori, T., Nazian, S. J., Tan, J., Potter, H. and Arendash, G. W. (2007). Enhanced cognitive activity—over and above social or physical activity—is required to protect Alzheimer’s mice against cognitive impairment, reduce Aβ deposition, and increase synaptic immunoreactivity. Neurobiology of learning and memory 88, 277–294.

Feehley, T., Plunkett, C. H., Bao, R., Hong, S. M. C., Culleen, E., Belda-Ferre, P., Campbell, E., Aitoro, R., Nocerino, R. and Paparo, L. (2019). Healthy infants harbor intestinal bacteria that protect against food allergy. Nature medicine 25, 448.

Gajbhiye, K. R., Gajbhiye, V., Siddiqui, I. A., Pilla, S. and Soni, V. (2017). Ascorbic acid tethered polymeric nanoparticles enable efficient brain delivery of galantamine: An in vitro-in vivo study. Scientific reports 7, 11086.

Halekoh, U. and Højsgaard, S. (2014). A Kenward-Roger approximation and parametric bootstrap methods for tests in linear mixed models - the R package pbkrtest. Journal of Statistical Software 59, 1–30.

Kuznetsova, A., Brockhoff, P. B. and Christensen, R. H. (2017). lmerTest package: tests in linear mixed effects models. Journal of Statistical Software 82, 1–26.

Michener, W. K. (2015). Ten simple rules for creating a good data management plan. PLoS Comput Biol 11, e1004525.

R Core Team. (2020). R: A Language and Environment for Statistical Computing. Vienna, Austria: R Foundation for Statistical Computing.

Ridaura, V. K., Faith, J. J., Rey, F. E., Cheng, J., Duncan, A. E., Kau, A. L., Griffin, N. W., Lombard, V., Henrissat, B. and Bain, J. R. (2013). Cultured gut microbiota from twins discordant for obesity modulate adiposity and metabolic phenotypes in mice. Science (New York, NY) 341.

Rosenzweig, N., Dvir-Szternfeld, R., Tsitsou-Kampeli, A., Keren-Shaul, H., Ben-Yehuda, H., Weill-Raynal, P., Cahalon, L., Kertser, A., Baruch, K. and Amit, I. (2019). PD-1/PD-L1 checkpoint blockade harnesses monocyte-derived macrophages to combat cognitive impairment in a tauopathy mouse model. Nature communications 10, 465.

Simpson, S. H. (2015). Creating a data analysis plan: What to consider when choosing statistics for a study. The Canadian Journal of Hospital Pharmacy 68, 311.

Tanoue, T., Morita, S., Plichta, D. R., Skelly, A. N., Suda, W., Sugiura, Y., Narushima, S., Vlamakis, H., Motoo, I. and Sugita, K. (2019). A defined commensal consortium elicits CD8 T cells and anti-cancer immunity. Nature 565, 600.

Venables, W. N. and Ripley, B. D. (2002). Modern Applied Statistics with S. New York: Springer.

Verbeke, G. and Molenberghs, G. (2000). Linear Mixed Models for Longitudinal Data. New York, NY: Springer New York.

